# L-Phenylalanine acts as a broad-spectrum plant growth biostimulator for both dicotyledonous and monocotyledonous crops

**DOI:** 10.64898/2026.11.16.766952

**Authors:** Yigal Elad, Dalia Rav-David, Michal Oren-Shamir

## Abstract

L-phenylalanine (Phe), an aromatic amino acid known to induce plant pathogen resistance at concentrations >4 mM, also functions as a broad-spectrum plant growth biostimulator in the total absence of biotic stress. To evaluate its biostimulatory potential, drench and foliar spray applications of Phe (1-16 mM) were tested on dicotyledonous (cucumber, tomato, lettuce) and monocotyledonous (wheat) crops grown under commercial-like net-house conditions with full NPK fertigation (150 mg/ L N). Weekly applications significantly enhanced vegetative growth across all species, increasing plant height by ≥22-26%, node numbers by ≥23-40%, and leaf/leaflet area by ≥36-48% in cucumber and tomato, as well as expanding leaf area in lettuce. In reproductive stages, Phe treatments increased flower initiation, set fruits (≥23-27%), total fruit number (≥31%), and fruit yield per plant (≥30-36%) in cucumber and tomato. In wheat, Phe increased tiller formation, total spike counts by ≥36%, and fresh/dry plant biomass. Growth enhancement required a strict concentration threshold of ≥4 mM; lower concentrations (1-2 mM) were inconsistent, and applying 2 mM twice weekly failed to replicate the significant gains of a single 4 mM weekly treatment. Because background fertigation provided a continuous and ample nitrogen supply, periodic trace applications of Phe contributed negligible amounts of nutritional N or C. This supports the conclusion that Phe acts through concentration-dependent physiological and metabolic biostimulation rather than nutrient supplementation.

## Introduction

Aromatic amino acids, including L-phenylalanine (Phe), are synthesized in plants via the shikimate pathway and serve as central precursors for secondary metabolites, flavonoids, and lignins essential for defense. Plants absorb externally given Phe from different organs very efficiently. Among else, Phe treatment results in increased flower fragrance and fruit aroma (Kumari et al., 2023). Furthermore, Exogenous pre-treatments with Phe at concentrations ≥4 mM reliably suppress a broad spectrum of fungal, bacterial, oomycetes, viral, and insect pests across diverse crops. During pathogen management trials, a striking secondary effect was observed: Phe-treated plants exhibited marked increases in vegetative vigor, canopy expansion, and harvest yield even in the complete absence of biotic stress (Elad et al., 2026; Kumar et al., 2024; Oliva et al., 2020; Patel et al., 2020. While observing the Phe effect on disease and pest in the various plant species we noticed a change in the plants growth and established new goals to our research – to test the effect of Phe on plant growth.

Improvement of plant growth by Phe was the subject of few publications. Foliar treatments with Phe concentrations of 100 and 200 mg/L that were applied to *Trigonella foenum-graecum* resulted in significant increase in growth parameters, yield components and antioxidant activity (Al-Duraid et al., 2019). Soil drench with Phe seedlings improved parameters of maize growth, such as height and fresh weight of shoots, distance between shoot nodes, and width of leaves (Sarwar and Frankenberger Jr, 1995). In poplar (*Populus × canescens*) trees the roots can uptake Phe as source of nitrogen which promotes the growth of the trees. The role of Phe as N source in this plant species is probably linked with ammonium ions re-utilization after the deamination of Phe (Jiao et al., 2018). Furthermore, previous studies leave several critical agronomic and biological questions unresolved: <u>Broad-spectrum efficacy</u>: Does exogenous Phe function universally across both dicotyledonous (cucumber, tomato, lettuce) and monocotyledonous (wheat) species?; <u>Application pathway</u> <u>equivalence</u>: Do root drench and foliar spray applications achieve comparable biostimulatory efficacy?; <u>Nutritional vs. metabolic triggering</u>: Does growth promotion stem from direct organic nitrogen (N) or carbon (C) supplementation, or does it represent a metabolic signaling trigger?; <u>Concentration thresholds</u>: Is biostimulation governed by a strict minimum concentration threshold (≥4 mM) or by cumulative dose over time?

To address these questions, this study systematically evaluated the biostimulatory effects of exogenous Phe across four major crop species (cucumber, tomato, lettuce, and wheat) grown under commercial-like net-house conditions. Importantly, all plants were continuously fertigated with complete NPK nutrient solution supplying 150 mg/L N (10.71 mM N), ensuring that growth responses to periodic trace applications of Phe reflect physiological and metabolic biostimulation rather than basic nutrient supply.

## Materials and methods

### Plant Materials, Growth Media, and Experimental Environment

Plants were grown in a horticultural mixture containing peat and tuff (7:3 v/v, unsorted to 10 mm). The four plant species were grown in experimental pest free net houses at the Volcani Institute, Rishon LeZion, Israel (31°5800900 N 34°4800200 E). Temperatures during the experimental plants’ growth were between 19 and 27°C, and RH was above 65%. Daylight duration was 10 to 14 h. (Elad et al. 2010, 2026, Graber et al. 2010, Gupta et al. 2021).

Cucumber (*Cucumis sativus* L.) cultivars used in experiments were Ilan (Zeraim Gedera, Israel) and Manny and Romi (Genesis Seeds, Israel). Cv. Ilan seeds were used for seedling experiments grown in 1 L pots after transplanting bare root 4 d old seedlings from a peat nursery pot and grown for two weeks as single plant in each pot. Cvs Manny and Romi were used for mature plants’ experiments that were grown as single plant in 10 L pots. The plants were grown under commercial- like conditions, initially without experimental treatments for 3 weeks and then treated for 4 to 5 weeks with the selected treatments, arranged in blocks with six potted-plant replicates. To prevent *Pythium* spp. damping off the nursery was treated by drenching once with a fungicide (0.25% Dynon in water, containing 722 g/L Propamocarb HCl, Bayer, Germany).

Two cultivars of tomato (*Solanum Lycopersicum* L.) were used: cv. Ikram (an indeterminate tomato type of Zeraim Gedera and Syngenta Vegetable Seeds, Israel) and cv. Brigade (a processing tomato determinate type of Asgrow Seed Company (operated by Bayer Crop Science, USA). The plants were seeded by Hishtil Nurseries (Ashkelon, Israel) and transplanted 4-6 weeks later in our experimental structures at the Volcani Institute. Tomato plants were grown for 2 weeks after transplanting to 10 L pots in a pest free structure and then maintained under commercial like conditions for 7 weeks. The tomato plants, one in each pot, were used in experiments arranged in blocks with 5 replicates.

Lettuce (*Lactuca sativa*) cv. Raviv (Hazera Seeds, Shikmim, Israel) was grown from seeds for 6 weeks in the nursery (Hishtil, Nurseries, Kefar Yedidia, Israel) and then transplanted into 1 L pots using seedling plugs. Two weeks after transplanting the leaves were evaluated. The plants (single lettuce plant in a pot) were used in experiments arranged in blocks with 7 replicates.

Wheat (*Triticum aestivum* L.) cv. Bet Hashita (Hazera (1939) Ltd. Israel) was seeded in 5 L pots. Six seeds were sown per pot and after 13 days, the seedlings were thinned, if needed, to standardized density of 5 plants per pot. The wheat plants were grown for 14 weeks until spikes yield was evaluated. The wheat pots, containing 5 plants each, were used in experiments arranged in blocks with 10 replicates.

### Fertigation and Water Management

All experimental plants received uniform, automated drip fertigation 2–3 times daily using a balanced 5:3:8 NPK fertilizer solution (Deshen Gat, Kiryat Gat, Israel). The solution supplied 150 mg L ¹ nitrogen (10.71 mM N), 38 mg/L phosphorus, and 185 mg/L potassium at an electrical conductivity (EC) of 2.2 dS/m, with irrigation adjusted to maintain ∼30% drainage. Continuous high fertigation ensured that background macro- and micronutrient demands were fully satisfied throughout the trial period. (Elad et al. 2010, 2026, Graber et al. 2010, Gupta et al. 2021).

### Phe treatments

Pure L-phenylalanine (Sigma Aldrich Israel Ltd.) was applied in final concentrations of 1 to 16 mM according to the specifics of the treatments of the experiments. The spray applications were performed using a 1 L hand sprayer that was fixed to emit fine droplets until full canopy coverage was obtained. The solution was applied at volume of 6-20 mL per each mature plant, depending on plant size. Drench applications consisted of 8 and 40 mL per 1 and 10 L pots, covering the volume of root zone with ∼5% drainage. Treatments in the young cucumber plants (cv. Ilan) were carried out 2 times, in 4 days intervals. In mature cucumber plants (cv. Manny and Romi), weekly applications were conducted until one week before the last evaluation date. Tomato plants were treated once every week except for one experimental set up where the plants were treated twice a week as mentioned in the results section. Lettuce leaves were initially treated 20 days after transplanting (day 0) and 3 days later (day 3) with 4- and 8-mM spray or drench. Wheat experiments were drench and spray treated with 2- and 4-mM Phe, once a week until one week before last evaluation time. Treatment with up to 32 mM Phe did not result in any phytotoxic effect on all the treated plants.

### Measured parameters of growth

- Cucumber young plants: Leaf area calculated from measures of width and length of the leaf blades;

- Cucumber mature plants: Vegetative growth parameters of above ground plant height, number of nodes, leaf area calculated from measures of width and length of the leaf blades, and reproduction parameters including the number of set fruits per plant, number of mature fruits in each plant, weight of fruits in each plant and the calculated average weight of a single fruit.

Tomato plants’ vegetative growth parameters were above ground plant height, number of nodes, leaf length and area of terminal leaflet calculated from measures of width and length of the leaflet. Reproduction parameters included the number of set fruits and flowers per inflorescent and per plant, number of mature fruits in each plant, weight of fruits in each plant and the calculated average weight of a single fruit. In one experimental setup the fresh and dry weight of the canopy was also measured at the end of the experiment.

Lettuce leaf area at days 11 and 16 was calculated using two perpendicular diameters.

Growth parameters in wheat included the number of spikes in each pot (5 plants), number of tillers formed per pot, and fresh and dry biomass weight per plant.

All vegetative and reproductive parameters evaluated across the four crop species are defined in Table 1. In parameters of vegetative growth in cucumber, tomato and lettuce, the measures of each parameter at two evaluation dates were used to calculate the percent increase in measures of the plant organs, expressed as percent change. The % change between the two evaluation dates was calculated according to the formula:

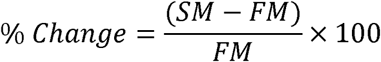

**Table 1.** Summary of experimental parameters, measurement methodologies, and evaluation timeframes across all four crop species.

| Crop | Target organ / stage | Evaluated parameter | Unit | Measurement method and timeframe | Figure in results |
|---|---|---|---|---|---|
| Cucumber |  |  |  |  |  |
|  | Seedlings | 1<sup>st</sup> true leaf | mm² | Calculated from length × width measurements at day 13 | Fig. S1 |
|  | Seedlings | Leaf area (leaves 1 and 2) | mm² | Calculated from length × width measurements at days 6 & 10 | Fig. 1b |
|  | Seedlings | Plant height | cm | Distance from root collar to apical meristem at day 10 | Fig. 1a |
|  | Vegetative | Plant height and node count | % change | % increase between days 4 and 19 | Figs 2ab |
|  | Vegetative | Leaf area (Leaf 10 / Leaf 14) | % change | Increase calculated from blade dimensions between days 0 and 10 | Fig. 2c |
|  | Vegetative | Nodes count | % change | Increase calculated between days 0 to 16 | Fig. S2a |
|  | Vegetative | Area of leaf 8 | % change | Increase calculated between days 0 to 7 | Fig. S2b |
|  | Vegetative | Area of leaf 14 | % change | Increase calculated between days 6 to 12 | Fig. S2c |
|  | Reproductive | Amount of set fruits | Number/plant | Day 16 | Fig. S2d |
|  | Reproductive | Number of mature fruits and weight of mature fruits | Number/plant, g/plant | 28 Days | Figs S2ef |
|  | Reproductive | Set fruits and mature fruits | No./plant | Total fruit counts recorded at days 19 and 28 post-treatment initiation | Figs 3ab |
|  | Reproductive | Fruit yield and single fruit weight | g/plant, g/ fruit | Total cumulative fruit mass and average individual fruit weight at day 28 | Figs 3cd |
|  | Vegetative | Plant height | Cm | Distance from root collar to apical meristem at day 43 | Fig S3a |
|  | Vegetative | Node count | % Change | Increase calculated between days 5 to 43 | Fig S3b |
|  | Vegetative | Leaf 12 area | % Change | Increase calculated between days 5 to 43 | Fig S3c |
|  | Reproductive | Set fruits | No/plant | Counted at day 50 | Fig S3d |
|  | Reproductive | Mature fruits | No/plant | Counted at day 56 | Fig S3e |
|  | Reproductive | Fruit yield and single fruit weight | g/plant, g/ fruit | Total cumulative fruit mass and average individual fruit weight at day 56 | Figs S3fg |
| Tomato | Vegetative | Plant height and node count | % change | % increase calculated between days 1 and 21 (or days 18 to 39) | Figs 4ab |
|  | Vegetative | Leaf length and terminal leaflet area | % change | Measured on leaf 5 and leaf 11 between days 1 and 21 | Figs 4cd |
|  | Reproductive | Set fruits and flowers, no. of fruits | No/inflorescence, No/plant | Counted at day 30 | Fig. 5abc |
|  | Reproductive | Fruit yield, Fruit number and single fruit weight | g/plant, no/plant, g/ fruit | Measured at day 56 | Fig 5def |
|  | Vegetative | Plant height and node count | % change | % increase calculated between days 18 to 39 | Table 1 Figs S4ab |
|  | Vegetative | Leaf length | % change | % increase calculated for leaf 5 between days 3 to 10, and leaf 11 between days 27 to 39 | Table 1 Figs S4cd |
|  | Vegetative | Terminal leaflet area | % change | % increase calculated for leaf 5 between days 3 to 10, and leaf 11 between days 27 to 39 | Table 1 Figs S4ef |
|  | Vegetative | Canopy biomass | g/plant | Wet and dry canopy mass recorded at harvest (Day 73) | Table 1 Figs S4gh |
|  | Reproductive | Inflorescences, set fruits | No/plant | Counted at days 27 and 39, respectively | Figs S5ij |
|  | Reproductive | Fruit yield, Fruit number and single fruit weight | g/plant, no/plant, g/ fruit | Measured day 73 | Figs S5klm |
| Lettuce | Vegetative | Canopy leaf area | % change | % increase calculated from two perpendicular leaf diameters between days 11 and 16 | Fig 6 |
| Wheat | Vegetative | Tiller count | No./pot | Total tillers per 5-plant pot evaluated at day 63 post-seeding | Fig 7d |
|  | Vegetative | Canopy biomass | g/plant | Whole-plant fresh and dry biomass evaluated at day 100 post-seeding | Figs 7 bc |
|  | Reproductive | Spike count | No./pot | Total productive spikes per 5-plants' pot evaluated at days 60 and 100 post-seeding | Fig 7a |

**Table 2.**
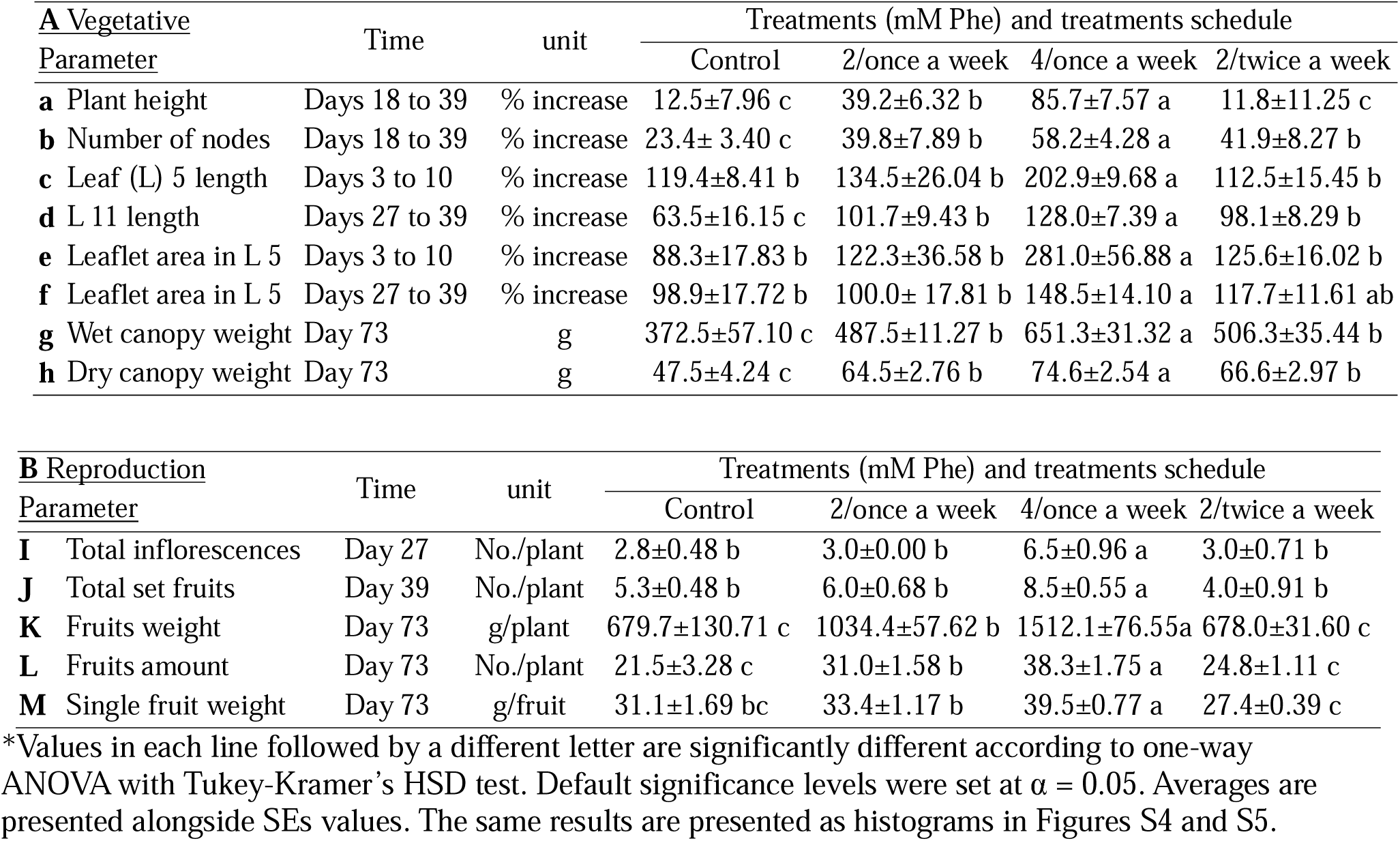
Effect of spray once and twice a week of 2 mM Phe and of 4 mM Phe sprayed once a week on growth of tomato plants as measured for vegetative (A) and production (B) parameters.

| **A** Vegetative | Time | unit | Treatments (mM Phe) and treatments schedule |  |  |  |
|---|---|---|---|---|---|---|
| Parameter |  |  | Control | 2/once a week | 4/once a week | 2/twice a week |
| **a** Plant height | Days 18 to 39 | % increase | 12.5±7.96 c | 39.2±6.32 b | 85.7±7.57 a | 11.8±11.25 c |
| **b** Number of nodes | Days 18 to 39 | % increase | 23.4± 3.40 c | 39.8±7.89 b | 58.2±4.28 a | 41.9±8.27 b |
| **c** Leaf (L) 5 length | Days 3 to 10 | % increase | 119.4±8.41 b | 134.5±26.04 b | 202.9±9.68 a | 112.5±15.45 b |
| **d** L 11 length | Days 27 to 39 | % increase | 63.5±16.15 c | 101.7±9.43 b | 128.0±7.39 a | 98.1±8.29 b |
| **e** Leaflet area in L 5 | Days 3 to 10 | % increase | 88.3±17.83 b | 122.3±36.58 b | 281.0±56.88 a | 125.6±16.02 b |
| **f** Leaflet area in L 5 | Days 27 to 39 | % increase | 98.9±17.72 b | 100.0± 17.81 b | 148.5±14.10 a | 117.7±11.61 ab |
| **g** Wet canopy weight | Day 73 | g | 372.5±57.10 c | 487.5±11.27 b | 651.3±31.32 a | 506.3±35.44 b |
| **h** Dry canopy weight | Day 73 | g | 47.5±4.24 c | 64.5±2.76 b | 74.6±2.54 a | 66.6±2.97 b |
| **B** Reproduction | Time | unit | Treatments (mM Phe) and treatments schedule |  |  |  |
| Parameter |  |  | Control | 2/once a week | 4/once a week | 2/twice a week |
| **I** Total inflorescences | Day 27 | No./plant | 2.8±0.48 b | 3.0±0.00 b | 6.5±0.96 a | 3.0±0.71 b |
| **J** Total set fruits | Day 39 | No./plant | 5.3±0.48 b | 6.0±0.68 b | 8.5±0.55 a | 4.0±0.91 b |
| **K** Fruits weight | Day 73 | g/plant | 679.7±130.71 c | 1034.4±57.62 b | 1512.1±76.55a | 678.0±31.60 c |
| **L** Fruits amount | Day 73 | No./plant | 21.5±3.28 c | 31.0±1.58 b | 38.3±1.75 a | 24.8±1.11 c |
| **M** Single fruit weight | Day 73 | g/fruit | 31.1±1.69 bc | 33.4±1.17 b | 39.5±0.77 a | 27.4±0.39 c |

\*Values in each line followed by a different letter are significantly different according to one-way ANOVA with Tukey-Kramer’s HSD test. Default significance levels were set at α = 0.05. Averages are presented alongside SEs values. The same results are presented as histograms in Figures S4 and S5.

Where FM is the first evaluation value, and SM is the second evaluation value.

### Data analysis

Experiments included 5 to 10 biological replicates for each treatment, as mentioned above specifically for each plant species. All experiments were repeated. Data in percentages were arcsine-transformed before further analysis. Standard errors (SE) of the means were calculated considering the degrees of freedom (DF = n - 1). As described in previous publications (Elad et al. 2010; 2017; 2021ab; Rav David et al. 2019). The data were analyzed using ANOVA and the Tukey-Kramer HSD test. Statistical analysis was performed (α = 5%) using JMP software (SAS Institute, Cary, NC, USA). Values of each parameter, graph or date followed by a different letter are significantly different from each other.

## Results

### Promotion of cucumber growth by Phe drench and spray Effect of Phe on cucumber seedlings

Cucumber seedlings were treated twice by Phe drench and spray, 2- and 4-mM (Figs 1 and S1). Dynon that is a fungicide aimed at *Pythium* spp. damping off was also used because it is a common treatment in cucumber nurseries, and we wished to verify the net effect of this fungicide. Size of the first true leaf was significantly increased by ≥46% by 4 mM Phe drench and by 2- and 4-mM Phe spray (Fig. 1a). Similarly, the size of the second true leaf was significantly increased by ≥13% by 4 mM Phe drench and by 2- and 4-mM Phe spray (Fig. S1). Plant height was significantly increased by ≥10% by 4 mM Phe drench and by 2- and 4-mM Phe spray (Fig. 1b). Phe 2 mM drench and dynon (Propamocarb HCl) drench did not significantly affect any of the 3 seedling growth parameters (Figs 1 and S1).

**Figure 1.**
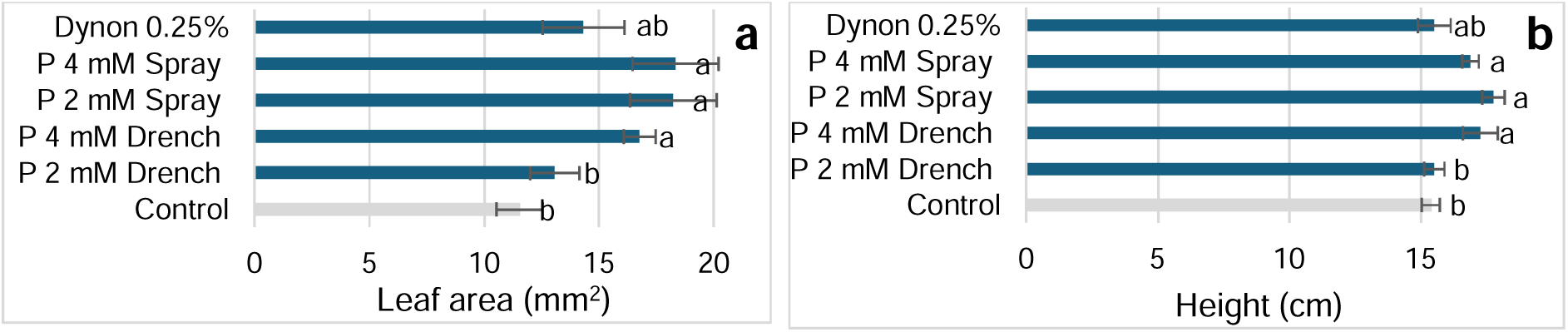
Growth of cv. Ilan cucumber seedlings as affected by drench and spray of 2- and 4-mM Phe (P) applied at 4 and 8 days after seeding and as affected by drench of dynon (Propamocarb HCl, Bayer, Germany) applied at the day of seeding. Effect of the treatments on the area of the first true leaf at 6 days (**a**), and on plants height at day 10 (**b**). Values in each graph followed by a different letter are significantly different according to one-way ANOVA with Tukey-Kramer’s HSD test. Default significance levels were set at α = 0.05. Bars = SEs.

### Effect of Phe on mature plants of cucumber

The growth of cucumber plants was followed until fruit harvest following weekly treatments of 4-, 8- and 16-mM Phe drench and spray (Figs 2 and 3). The change in plant height was significantly higher by ≥22% as a result of all Phe treatments, compared with the untreated control (Fig. 2a). The change in number of nodes was significantly higher by ≥23% as a result of all Phe treatments compared with the control (Fig. 2b). The change in leaf area was significantly higher by ≥48% as a result of 4- and 8-mM Phe treatments compared with the control (Fig. 2c).

**Figure 2.**
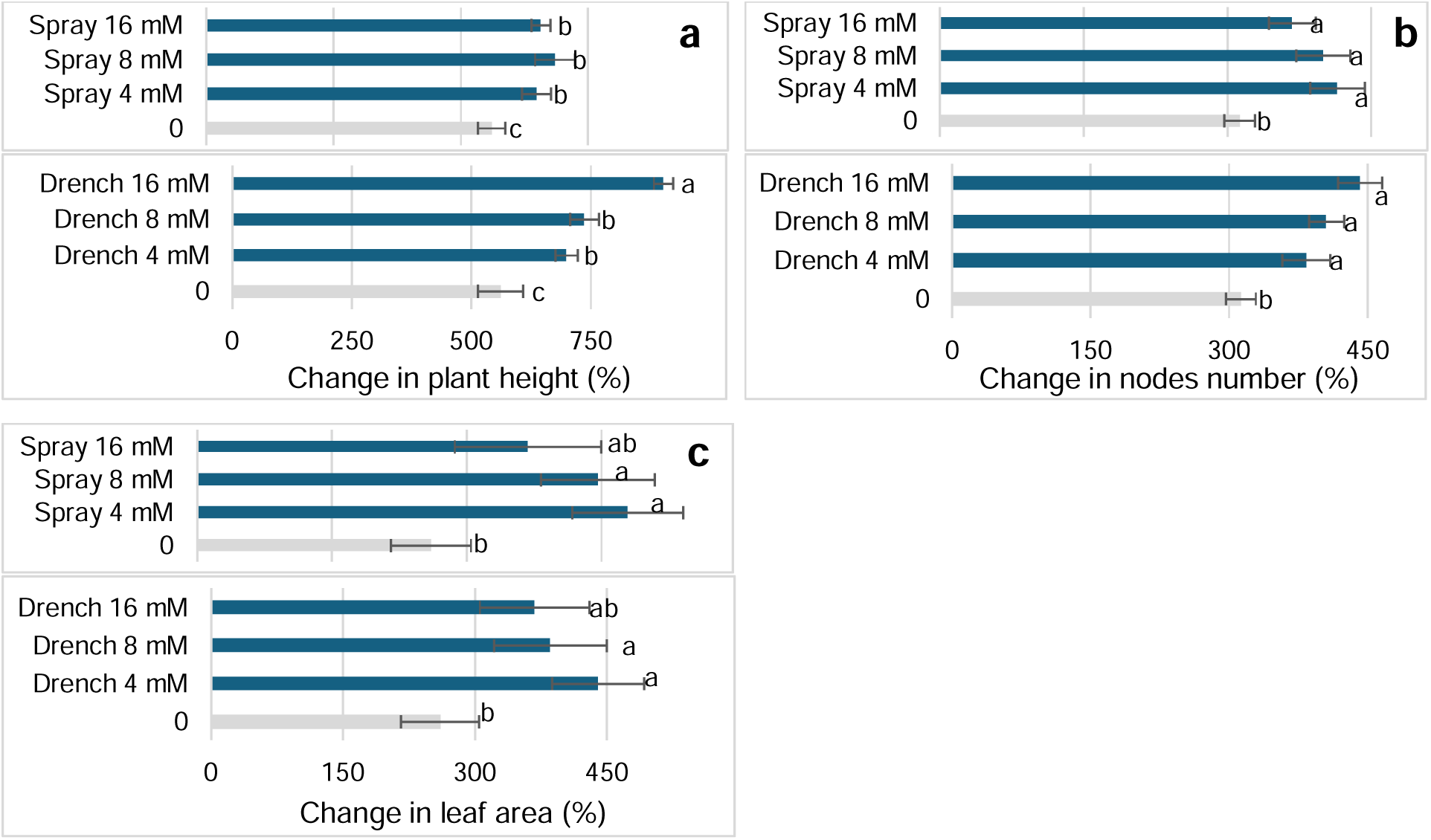
Effect of drench and spray of 4-, 8-, and 16-mM Phe on growth of cucumber plants cv. Manny grown in large pots and treated once a week for 3 weeks. The vegetative growth parameters that were evaluated were: **a**. plant height (change between days 4 to 19), **b**. nodes number (change between days 4 to 19, and **c**. area of leaf 10 (change between days 0 to 10). Values in each parameter followed by a different letter are significantly different according to one-way ANOVA with Tukey- Kramer’s HSD test. Default significance levels were set at α = 0.05. Bars = SEs. Note that the control column in the two graphs of each parameter represents the same treatment.

**Figure 3.**
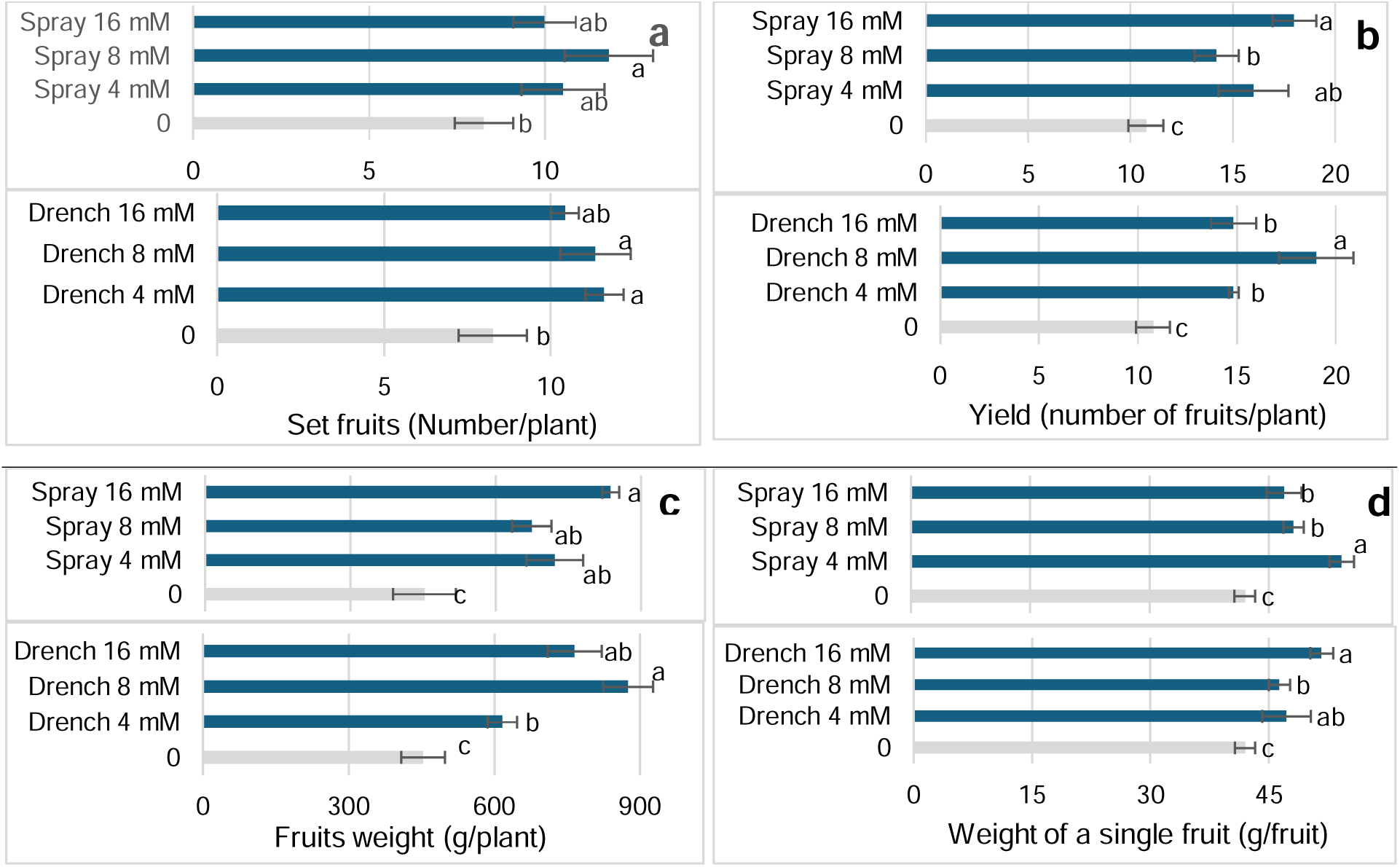
Effect of drench and spray of 4-, 8-, and 16-mM Phe on growth of cucumber plants (as detailed in Fig. S2). The reproductive growth parameters that were evaluated were: **a.** the number of set fruits at day 19, **b.** the number of mature fruits at day 28, **c.** weight of mature fruits at day 28, and **d.** weight of single fruits at 28 days. Values in each parameter followed by a different letter are significantly different according to one-way ANOVA with Tukey-Kramer’s HSD test. Default significance levels were set at α = 0.05. Bars = SEs. Note that the control column in the two graphs of each parameter represent the same treatment.

As for the reproduction parameters (Fig. 3), the number of set fruits was significantly increased by ≥27% with 4- and 8-mM Phe drench and 8 mM Phe spray as compared with the untreated control (Fig. 3d). The number of fruits increased by ≥31% (Fig. 3e) and the total weight of fruits was increased by ≥36% (Fig. 3f) by all Phe treatments as compared with the control. The average weight of a single fruit was significantly higher by ≥11% as a result of all Phe treatments compared with the untreated control (Fig. 3g).

Growth of cucumber plants was also followed with weekly treatments of 1-, 2-, 4-, 8-, and 16- mM Phe drench and spray (Fig. S2). The change in the number of nodes was significantly higher by ≥16% as a result of drench with 4 to 16 mM, and spray with 2 to 16 mM Phe as compared with the untreated control (Fig. S2a). The change in the area of leaf no. 8 was significantly higher by ≥34% as a result of 4 to 16 mM Phe drenches, and 2 to 16 mM Phe sprays, as compared with the control (Fig. S2b). The change in the area of leaf no. 14 was higher by ≥74% with all the Phe treatments, as compared with the control (Fig. S2c). The number of set fruits was significantly increased by ≥19% with all the Phe treatments as compared with the untreated control (Fig. S2d). The number of fruits was increased by ≥19% with treatments of drench of 4 to 16 mM Phe and with spraying of 2 to 16 mM Phe (Fig. S2e). The total weight of fruits was increased by ≥24% with treatments of drench of 4 to 16 mM Phe and with spraying of 2 to 16 mM Phe (Fig. S2f).

In order to test weather doubling the number of spray applications of 2 mM Phe from one application per week to twice every week we grew cucumber plants cv. Romi, treated them for 6 weeks and collected growth parameters (Fig. S3). A 4 mM spray treatment significantly increased cucumber growth compared to all other treatments as measured for plant height at day 43 (Fig. S3a), change in nodes number between days 5 and 43 (Fig. S3b), increase in the area of leaf 12 between days 12 and 19 (Fig. S3c), number of set fruits at day 50 (Fig. S3d), number, general weight of fruits and the average weight of a single fruit until week 8 (Fig. S3e-g). Results of the 2 mM Phe spray once a week and twice a week were similar to the control (Fig. S3).

### Phe promotes the vegetative growth and the reproductive stage of tomato

#### Effect of Phe on parameters of tomato vegetative growth

Tomato plants treated on a weekly basis for 7 weeks by spray and drench of Phe at concentrations between 1 to 8 mM. The height of plants significantly increased by ≥26% with drench of 1 to 4 mM Phe and by spray of 2 to 4 mM Phe (Fig. 4a). The number of nodes significantly increased by 40% with 2- and 4-mM drench and with 2 to 8 mM spray (Fig. 4b). The length of leaf 5 was significantly increased by ≥140% with ≥2 mM Phe drench and with 4- and 8-mM Phe spray (Fig. 4c). The area of leaflet in leaf 5 was significantly increased by ≥36% with drench treatments 1 to 8 mM Phe and with 2 to 8 mM Phe spray (Fig. 4d).

**Figure 4.**
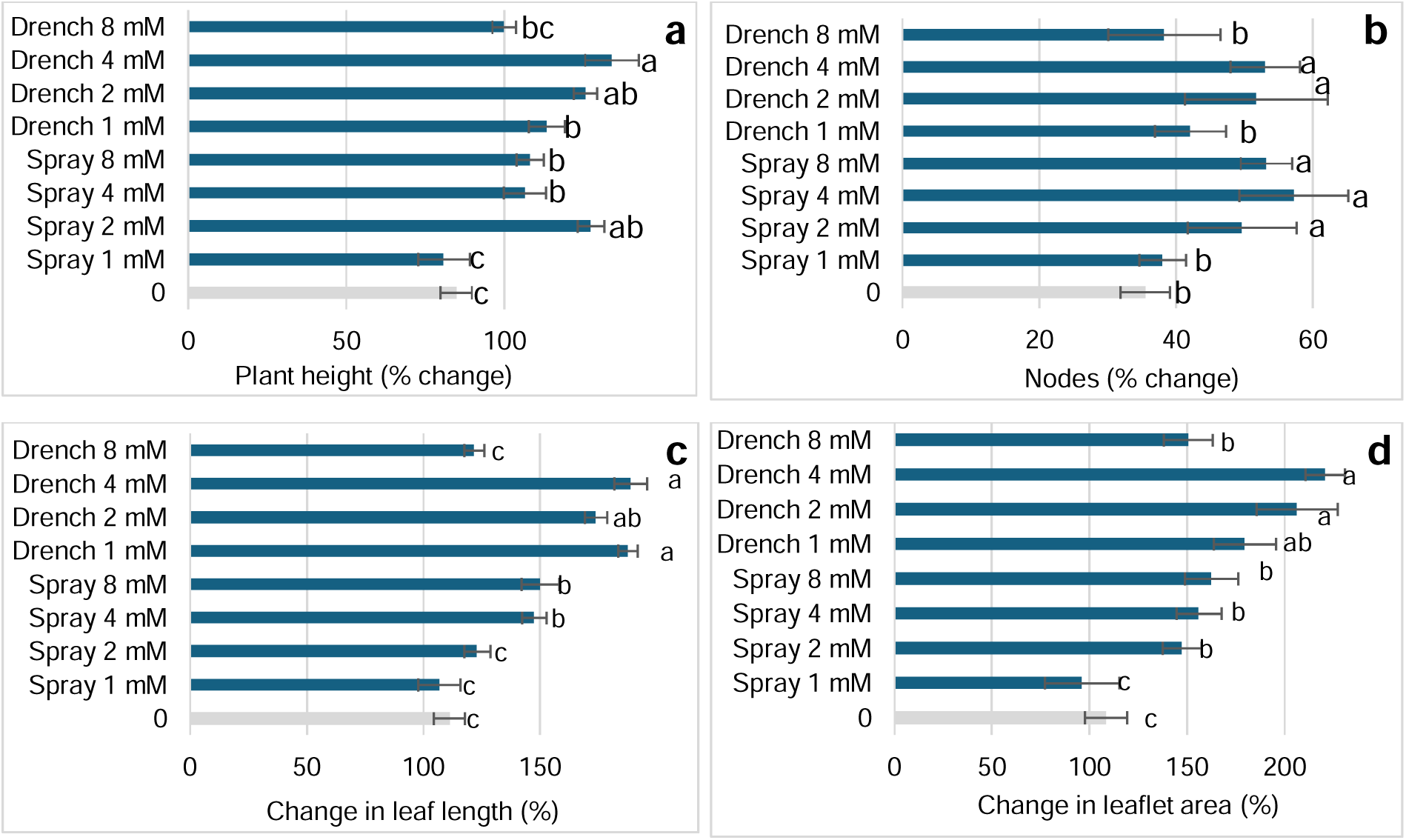
Effect of drench and spray of Phe in concentrations of 1 to 8 mM on vegetative growth parameters of plants of tomato cv. Ikram grown under commercial like conditions. Phe treatments were carried out once a week until day 42 after the initiation of applications. The parameters evaluated were **a**. change in plant height from day 1 to day 21; **b**. change in the number of nodes from day 1 to day 21; **c**. change in length of leaf 5 from day 1 to day 21; **d**. change in the terminal leaflet area (at leaf 5) from day 1 to day 14. Values in each parameter followed by a different letter are significantly different according to one-way ANOVA with Tukey-Kramer’s HSD test. Default significance levels were set at α = 0.05. Bars = SEs.

### Effect of Phe on reproductive parameters in tomato plants

The average number of fruits and flowers per inflorescence was increased by ≥34% with 2 to 8 mM drench and with 4- and 8-mM Phe spray (Fig. 5a). The total number of flowers and set fruits was significantly increased by ≥23% with 2- and 4-mM Phe drench and with 1 to 4 mM Phe spray (Fig. 5b). The total number of fruits at day 30 was significantly increased by ≥143% with drench of 2 to 8 mM and with spray of 2- and 4-mM Phe (Fig. 5c). Total number of fruits at day 46 was significantly increased by ≥61% only with drench 4 mM and 4 mM Phe spray (Fig. 5d). Total fruit weight at day 46 was significantly increased by ≥30% with drench of 2 to 8 mM Phe and with spray of 2 to 8 mM Phe (Fig. 5e). The calculated single fruit weight at day 46 was significantly increased by ≥13% with 8 mM Phe drench and 1 to 8 mM Phe spray (Fig. 5f).

**Figure 5.**
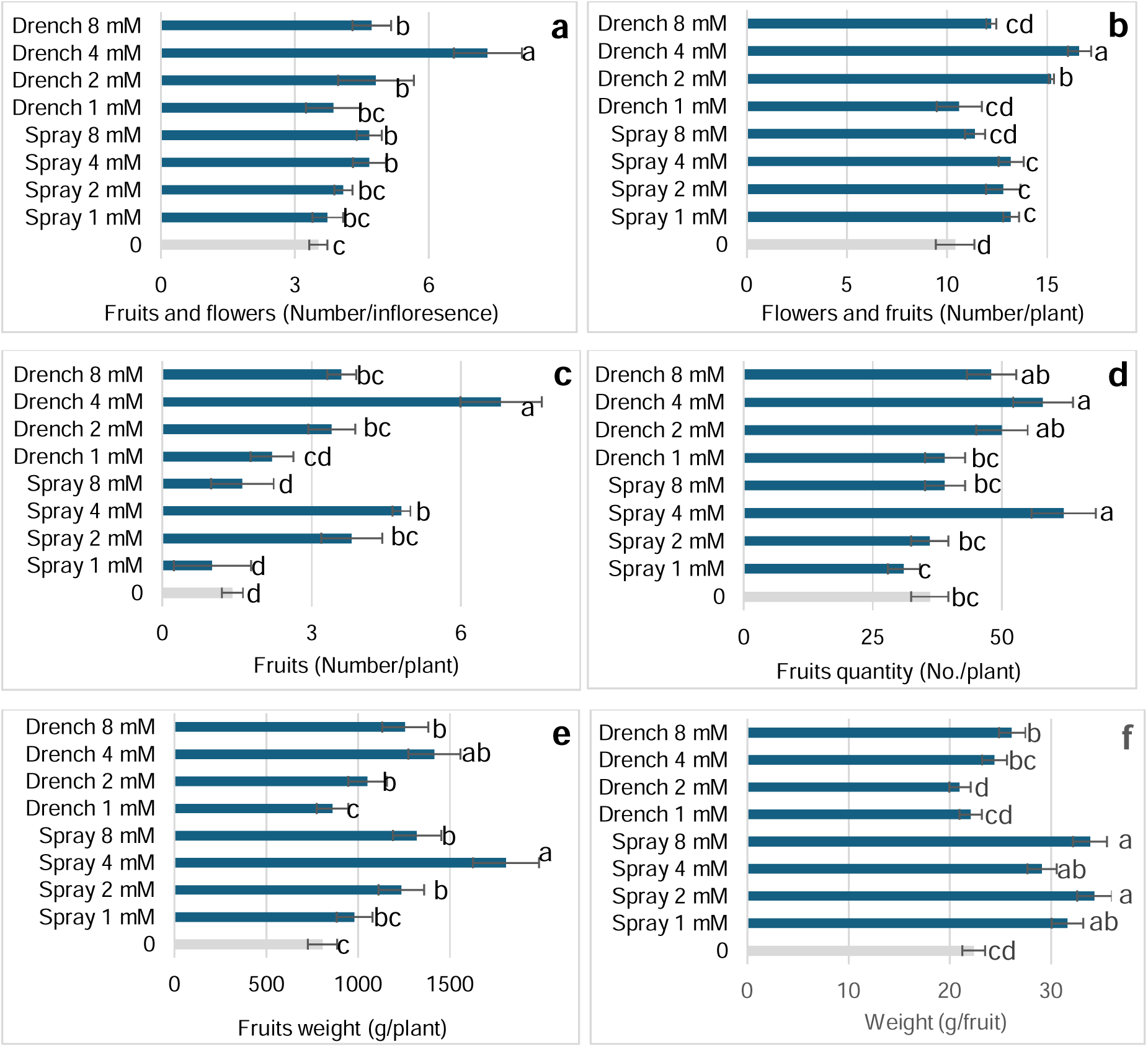
Effect of drench and spray of Phe in concentrations of 1 to 8 mM on reproductive parameters of tomato plants cv. Ikram grown under commercial like conditions. **a**. average number of set fruits and flowers at day 30; **b**. total number of flowers and fruits at day 30; **c**. number of fruits at day 30; **d**. fruit weight at day 46; **e**. fruits number at day 46; **f**. average weight of single fruit at day 46. Values in each parameter followed by a different letter are significantly different according to one-way ANOVA with Tukey-Kramer’s HSD test. Default significance levels were set at α = 0.05. Bars = SEs.

### Spray application of 2 mM Phe once and twice a week evaluated with vegetative growth and reproduction parameters in tomato

We tested if twice a week 2 mM spray application can induce a similar effect to 4 mM Phe applied once a week as carried out commonly in our experiments. Vegetative growth of tomato plants and production of flowers and fruits were evaluated during and 9 weeks of Phe spray applications of 4 mM once a week and 2 mM once and twice a week (Table 1 AB and Fig. S5). Phe spray of 4 mM significantly increased the values of vegetative growth as measured for plant height, nodes number, leaves length, leaflets area and canopy weight and also the flower and fruit production and plant fresh and dry weight (Table 1A). Spray with 2 mM Phe once a week was significantly different from the untreated control in evaluations of plant height, nodes number, change in leaf 11 length, total fruit weight, and number of fruits (Table 1B). Nevertheless, the twice-a-week spray schedule was in no measurement significantly better than the once-a-week 2 mM sprays that by itself was generally inferior to the 4 mM Phe.

### Phe promotes lettuce growth

Leaf area was significantly increased by 4- and 8-mM Phe spray, as well as by 8 mM Phe drench (Fig. 6).

**Figure 6.**
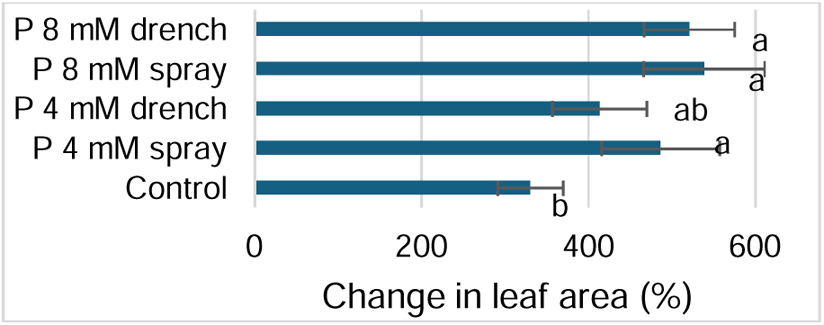
Effect of Phe (4- and 8-mM) applied by spray or drench in days 0 and 3 on lettuce leaf area. Increase in leaf area was calculated between leaf area measurements taken at days 11 and 16. Different letters indicate significant differences (one-way ANOVA with Tukey-Kramer HSD, α = 0.05). Error bars represent SE.

### Promotion of wheat growth by Phe

Phe was applied by spray and drench starting two weeks after seeding the wheat (Fig. 7). Spike number was assessed at 60 and 100 days after seeding, with a significant increase observed at day 100 for 2- and 4-mM Phe drench and 4 mM Phe spray which increased spike number by ≥36% (Fig. 7a). Whole plant fresh weight at day 100 was significantly increased by 2 mM Phe applied as drench and with 2- and 4-mM spray, and the dry weight of plants was increased significantly by the four tested treatments (Figs 7b,c). Tillers number at day 63 was increased by 4 mM Phe drench and spray (Fig. 7d); 2 mM treatments were not evaluated for this parameter.

**Figure 7.**
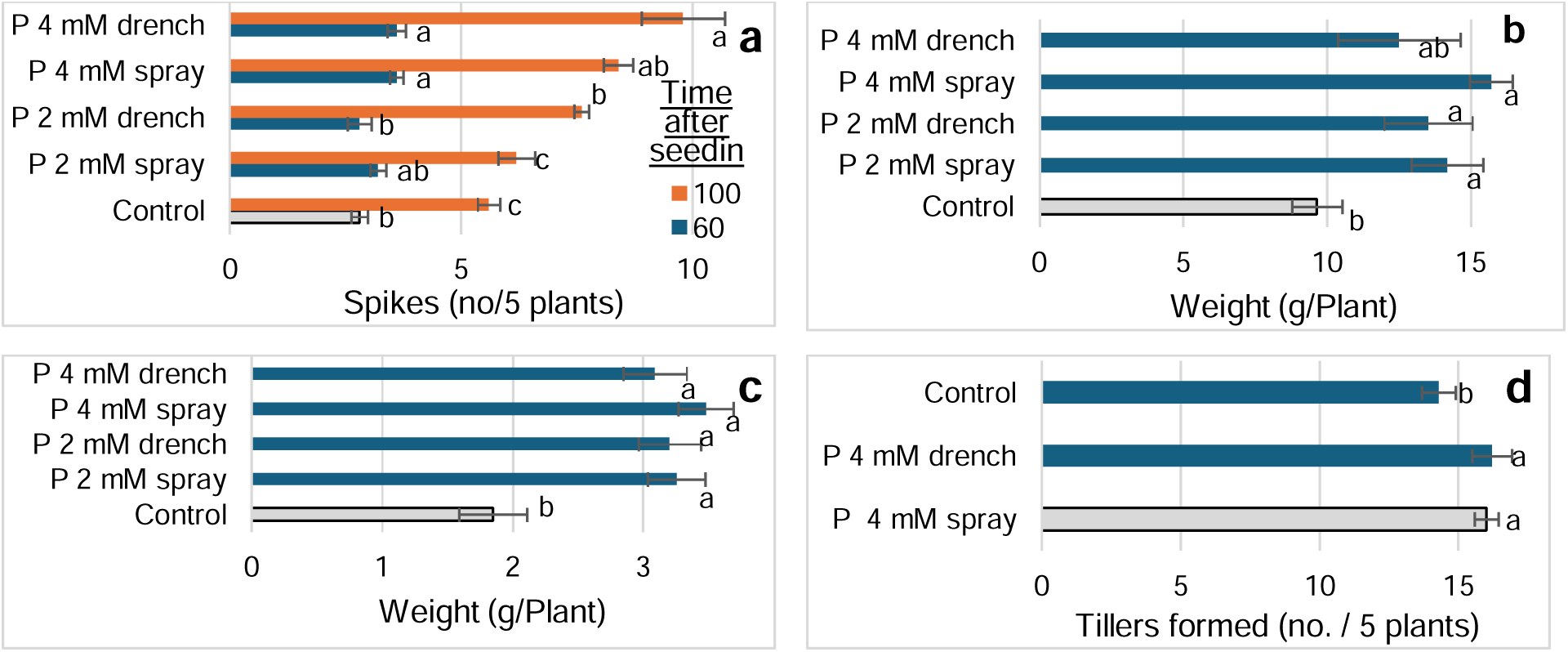
Effect of Phe on wheat growth. Phe (2- and 4-mM) was applied by spray and drench to cv. Bet Hashita starting at the two-leaf stage, with weekly applications until day 92 after seeding. Measured parameters included: **a.** spike number at days 60 and 100, **b.** fresh and **c.** dry biomass at day 100, and **d**. in a separate experiment, tillers number evaluated at day 63. Different letters indicate significant differences for each parameter and time point (one-way ANOVA with Tukey-Kramer HSD, α = 0.05). Error bars represent SE.

## Discussion

### Broad-spectrum biostimulatory efficacy across dicotyledonous and monocotyledonous crops

L-phenylalanine (Phe) that is applied externally not only contributes to disease control (Elad et al., 2026; Patel et al., 2020; Oliva et al., 2020) in plants but, as described here for four plant species, it also influences growth and developmental processes in these plants. Phe is involved in early seedling functions such as cell defense, cell division, and plastid development (Perkowski and Warpeha 2019), increased growth parameters, yield components and antioxidant activity in fenugreek (*Trigonella foenum-graecum*) (Al-Duraid et al., 2019), and here we also demonstrate effects in young as well as in mature plants, including cucumber (Figs 1, 2,3,S1,S2 and S3), tomato (Table 1, Figs 4, 5, S4 and S5) and lettuce (Fig. 6). There is limited information on the effect of Phe on the growth of these plant species. Combination of 25 ppm (=0.168 mM) methionine with 100 ppm (=0.605 mM) Phe improved cucumber biomass production; Foliar application of the Phe improved nutrients (Marium et al., 2021). Growth variables in tomato were not affected by the presence of the amino acids (AAs) alanine, serine, tyrosine and Phe, each at 0.2 mM. Yet, Phe and the other AAs in the tomato nutrient solution had a beneficial effect on the leaf mineral status and on the chlorophyll concentration of the leaves (Garcia et al., 2011). Similarly, aspartic acid with glutamic acid, increased the growth of tomato seedlings and L-alanine decreased it (Alfosea-Simón et al., 2021). For lettuce grown as nursery plugs in small cells, foliar spray with 11 different AAs (excluding Phe), each alone or combined, changed positively or negatively the lettuce fresh and dry weight and the leaf area (Salinas et al., 2019).

In the present research, effect of externally applied Phe was observed not only in dicots but also in a monocot plant species – wheat (Fig. 7). In wheat we observed increase of tillers formation, spikes number and general weight of the plant. There are no similar reports on the effect of Phe on wheat but in maize (*Zea mays*), soil drench application of Phe at concentration of up to 20 mg/kg soil had a significant effect on vegetative growth parameters in shoots and leaves and the authors concluded that the effect stemmed from increased phenylacetic acid that is a metabolic product derived from Phe (Sarwar & Frankenberger, 1995). Foliar application of 0.5 g/L glutamate and Phe (=3.40 and 3.03 mM, respectively) increased the seed-setting rate, grain yield, and the contents of chlorophyll *a*, chlorophyll *b*, and carotenoid in fragrant rice (Luo et al., 2023). Also, Phe, tryptophan and tyrosine at 0.5 to 2.0 g/L (=3.03 to 12.12 mM Phe) increased grain yield and seed-setting rate of fragrant rice (Lou et al., 2024).

### Concentration threshold kinetics overrule cumulative mass action

Throughout the current research the Phe was applied at various concentrations ≥1, 2, and 4 mM. Biostimulatory responses to exogenous Phe were strictly governed by a minimum concentration threshold of ≥4 mM. Across multiple trial sets, 1 mM and 2 mM applications produced inconsistent or non-significant growth responses.

This concentration dependency was further clarified by our dose-frequency trials in cucumber and tomato. Applying 2 mM Phe twice weekly supplied an identical cumulative chemical mass over time as 4 mM applied once weekly. Yet, 2 mM applied twice weekly failed to induce significant vegetative or reproductive gains, whereas a single weekly 4 mM application triggered maximum biostimulatory responses. This key outcome proves that Phe biostimulation operates via a concentration-dependent physiological trigger requiring ≥4 mM to activate downstream metabolic cascades, rather than a cumulative mass-action effect. Notably, this ≥4 mM threshold aligns directly with the concentration requirements reported for Phe-induced disease resistance and parallels concentration ranges documented in fragrant rice (≥3.03 mM) (Luo et al., 2023, 2024) and 0.605 mM Phe in cucumber (Marium et al., 2021). Phe can be applied on plants either by drench (Koca & Karaman 2015; Feng et al. 202) or spray (Bahadur et al. 2012; Luo et al., 2023; Marium et al., 2021) resulting in positive growth results. In the present research both Phe methods of application resulted in significantly the same results.

Growth promotion operates via metabolic signaling rather than direct N nutritive supplementation

Phe as an AA may affect plants’ growth as an organic nitrogen source and even as metabolic substrate (Jiao et al., 2018). However, multiple lines of evidence from our experimental design refute a direct nutritional mechanism: There was a background fertigation load in our experimental setups All experimental plants received continuous automated drip fertigation supplying 150 mg L ¹ N (10.71 mM N) 2–3 times daily. Weekly applications of Phe (delivered in small volumes of 6–20 mL spray or 8–40 mL drench per plant) provided a negligible fraction of N and C relative to the background fertigation regime. Therefore, there were only trace quantities of exogenous Phe that was supplied to the plants. Moreover, solitary weekly pulses of organic N would create nutrient fluctuations in the root zone and nutritional imbalance risks. Furthermore, productive fruit filling requires balanced potassium co-provision (typically at 1:1.5 to 1:2 N:K ratio); isolated N loading promotes purely vegetative luxus consumption rather than the concurrent fruit set and yield gains observed here. Consequently, the growth enhancements elicited by exogenous Phe represent genuine physiological and metabolic biostimulation rather than basic nutrient supplementation.

### Putative physiological and rhizosphere mechanisms

The equivalent performance of root drench and foliar spray applications indicates that Phe biostimulation can be triggered through distinct physiological entry points.

Root-applied Phe may further promote plant growth by modulating the nitrogen cycling and shaping the rhizosphere microbiome, increasing beneficial bacterial taxa while reducing harmful ones (Koca & Karaman 2015; Feng et al. 2023). Thus, root-applied Phe can reshape the root-zone microenvironment by recruiting beneficial nitrogen-cycling bacterial taxa while suppressing detrimental rhizosphere microbes. This microbial modulation enhances nutrient availability and root system efficiency under soilless substrate conditions.

Foliar application of Phe has been shown to increase phenylalanine ammonia-lyase (PAL) activity, as observed in pea leaves (Bahadur et al. 2012), which regulates lignin biosynthesis and a wide range of phenolic secondary metabolites. Accordingly, aromatic amino acids such as phenylalanine and tyrosine are key drivers of lignification pathways in plants, as demonstrated in wheat (Feduraev et al. 2020). Exogenous foliar application of Phe elevates the activity of PAL, the primary rate-limiting enzyme bridging primary metabolism to the phenylpropanoid pathway. Increased PAL activity promotes phenolic secondary metabolism and lignification. In cereal crops such as wheat, aromatic amino acids drive cell wall lignification and structural growth, while in maize, exogenous Phe is metabolized into phenylacetic acid (PAA), a naturally occurring auxin-type growth regulator that directly stimulates cell elongation and nodal development (Sarwar and Frankenberger Jr, 1995).

## Conclusion

This study demonstrates that exogenous L-phenylalanine (Phe) functions as a broad-spectrum, non- nutritional biostimulator capable of enhancing vegetative development and reproductive yield across diverse dicotyledonous (cucumber, tomato, lettuce) and monocotyledonous (wheat) crops under commercial net-house conditions. Applied via root drench or foliar spray, Phe biostimulation requires a strict concentration threshold of ≥4 mM and operates independently of cumulative mass- action or direct nitrogen fertilization. Integrating periodic Phe applications into commercial programs offers a robust, eco-friendly strategy to optimize crop productivity and yield efficiency. sed above.

## Supporting information

Supplemental Figures Elad et al

## ACKNOWLEDGEMENTS

We thank the late Eyal Cohen RIP, Dr. Galit Sharabani, and Ohad Zuckerman of Copia for the encouragement and advice during the PostBoost project development and for the support. We thank members of the Elad and Oren-Shamir research groups at the Volcani Institute for their continuous support. The fruitful discussions and help of Dr. Noam Alkan of the Institute of Postharvest and Food Sciences, Volcani Institute is much appreciated. We thank Menachem Borenshtein, Eyal Glanz of the Plant Pathology and Weed Research Department for the technical support in the greenhouses. The warm and interesting discussions and the excellent implementation of the following people of ICA International Chemicals (Pty) Ltd. South Africa is very much appreciated: Wouter Schreuder Jr., the late Kobus Serfontein RIP, Suzel Serfontein, Pieter Louw, and Gert van Coller.

## Author Contributions

**Yigal Elad**: Writing – review & editing, Writing – original draft, Supervision, Methodology, Investigation, Formal analysis, Conceptualization. **Dalia Rav-David**: Methodology, Investigation, Formal analysis. **Michal Oren Shamir**: Methodology, writing – review & editing, Supervision, Methodology, Investigation, Funding acquisition, Formal analysis, Conceptualization.

## Data Availability Statement

The authors declare that the data supporting the findings of this study are available within the paper and its Supporting Information files. Raw data is available from the corresponding author upon reasonable request.

