## Supplemental Figures Elad et al for "L-Phenylalanine acts as a broad-spectrum plant growth biostimulator for both dicotyledonous and monocotyledonous crops"

**Supplemental material**

Leaf area (mm2)

|  | 2nd leaf area 13d |
| --- | --- |
| Control | 120.92 |
| P 2 mM Drench | 129.18 |
| P 4 mM Drench | 136.44 |
| P 2 mM Spray | 143.344 |
| P 4 mM Spray | 141.71 |
| Dynon 0.25% | 121.512 |

Figure S1

Growth of cv. Ilan cucumber seedlings as affected by drench and spray of 2- and 4-mM Phe applied at 4 and 8 days after seeding and as affected by drench of dynon (Propamocarb HCl, Bayer, Germany) applied at the day of seeding. Effect of the treatments on the area of the first true leaf at day 13. Other results are described in Fig. 1. Values followed by a different letter are significantly different according to one-way ANOVA with Tukey–Kramer’s HSD test. Default significance levels were set at α = 0.05. Bars = SEs.

a

|  | 0-16 |
| --- | --- |
| 0 | 152.679 |
| Spray 1 mM | 170.238 |
| Spray 2 mM | 193.829 |
| Spray 4 mM | 224.802 |
| Spray 8 mM | 208.095 |
| Spray 16 mM | 300 |

b

|  | Leaf 8, 0-7 |
| --- | --- |
| 0 | 111.521 |
| Spray 1 mM | 143.485 |
| Spray 2 mM | 158.01 |
| Spray 4 mM | 194.903 |
| Spray 8 mM | 158.01 |
| Spray 16 mM | 146.151 |

Change in number of nodes (%)

|  | 0-16 |
| --- | --- |
| 0 | 152.679 |
| Drench 1 mM | 155.675 |
| Drench 2 mM | 163.333 |
| Drench 4 mM | 177.56 |
| Drench 8 mM | 191.944 |
| Drench 16 mM | 205.622 |

Change in leaf area (%)

|  | Leaf 8, 0-7 |
| --- | --- |
| 0 | 111.521 |
| Drench 1 mM | 125.923 |
| Drench 2 mM | 126.802 |
| Drench 4 mM | 148.855 |
| Drench 8 mM | 162.151 |
| Drench 16 mM | 172.073 |

c

|  | Leaf 14, 6-12 |
| --- | --- |
| 0 | 240.992 |
| Spray 1 mM | 545.412 |
| Spray 2 mM | 594.644 |
| Spray 4 mM | 460.675 |
| Spray 8 mM | 506.565 |
| Spray 16 mM | 486.05 |

d

|  | 16 d |
| --- | --- |
| 0 | 8.4 |
| Spray 1 mM | 8.66667 |
| Spray 2 mM | 9.83333 |
| Spray 4 mM | 13.3333 |
| Spray 8 mM | 12 |
| Spray 16 mM | 11.5 |

Change in leaf area (%)

|  | Leaf 14, 6-12 |
| --- | --- |
| 0 | 240.992 |
| Drench 1 mM | 442.974 |
| Drench 2 mM | 418.913 |
| Drench 4 mM | 492.863 |
| Drench 8 mM | 438.95 |
| Drench 16 mM | 531.131 |

Set fruits (number/plant)

|  | 16 d |
| --- | --- |
| 0 | 8.4 |
| Drench 1 mM | 10.4 |
| Drench 2 mM | 10.5 |
| Drench 4 mM | 10.7143 |
| Drench 8 mM | 12.3333 |
| Drench 16 mM | 15.3333 |

e

|  | Total 4 weeks |
| --- | --- |
| 0 | 10 |
| Spray 1 mM | 10.25 |
| Spray 2 mM | 14 |
| Spray 4 mM | 16 |
| Spray 8 mM | 14 |
| Spray 16 mM | 18.75 |

f

|  | Total 4 weeks |
| --- | --- |
| 0 | 516.314 |
| Spray 1 mM | 465.907 |
| Spray 2 mM | 723.44 |
| Spray 4 mM | 808.993 |
| Spray 8 mM | 775.632 |
| Spray 16 mM | 987.015 |

Fruit yield (number/plant)

|  | Total 4 weeks |
| --- | --- |
| 0 | 10 |
| Drench 1 mM | 8.6 |
| Drench 2 mM | 10.5 |
| Drench 4 mM | 11.8571 |
| Drench 8 mM | 14.6667 |
| Drench 16 mM | 13.8333 |

Fruit yield (g/plant)

|  | Total 4 weeks |
| --- | --- |
| 0 | 516.314 |
| Drench 1 mM | 451.828 |
| Drench 2 mM | 572.558 |
| Drench 4 mM | 642.23 |
| Drench 8 mM | 739.478 |
| Drench 16 mM | 814.372 |

Figure S2

Effect of drench and spray of 1-, 2-, 4-, 8-, and 16-mM Phe on the growth of cucumber plants cv. Manny grown in large pots and treated once a week for 3 weeks. The growth parameters that were evaluated were, **a.** nodes number (change between days 0 to 16), **b**. area of leaf 8 (change between days 0 to 7), **c**. area of leaf 14 (change between days 6 to 12), **d**. number of set fruits (day 16), **e**. number of mature fruits (28 days), and **f**. weight of mature fruits (28 days). Values in each parameter followed by a different letter are significantly different according to one-way ANOVA with Tukey–Kramer’s HSD test. Default significance levels were set at α = 0.05. Bars = SEs. Note that the control column in the two graphs of each parameter represent the same treatment.

a

Plant height (cm)

|  | 43 d |
| --- | --- |
| Control | 159.2 |
| 2 mM once/week | 139.4 |
| 4 mM once/week | 184 |
| 2 mM twice/week | 151.8 |

b

Change in nodes number (%)

|  | Percent change 5 to 43 |
| --- | --- |
| Control | 195.884 |
| 2 mM once/week | 96.1142 |
| 4 mM once/week | 248.056 |
| 2 mM twice/week | 198.333 |

c

Change in leaf area (% increase)

|  | Perc increase 12 to 19d |
| --- | --- |
| Control | 149.891 |
| 2 mM once/week | 248.444 |
| 4 mM once/week | 454.285 |
| 2 mM twice/week | 214.681 |

d

Set fruits (Number/plant)

|  | 50 |
| --- | --- |
| Control | 34.2 |
| 2 mM once/week | 30.2 |
| 4 mM once/week | 41.4 |
| 2 mM twice/week | 34.2 |

Fruits (number/plant)

|  | num |
| --- | --- |
| Control | 33.2 |
| 2 mM once/week | 29.2 |
| 4 mM once/week | 39.2 |
| 2 mM twice/week | 31.2 |

Fruits weight (g/plant)

|  | weight |
| --- | --- |
| Control | 1261.8 |
| 2 mM once/week | 1109.99 |
| 4 mM once/week | 1758.52 |
| 2 mM twice/week | 1197.54 |

g

Single fruit weight (g/fruit)

|  | fruit weight |
| --- | --- |
| Control | 38.1798 |
| 2 mM once/week | 37.8511 |
| 4 mM once/week | 44.9528 |
| 2 mM twice/week | 38.4014 |

Figure S3

Effect of spray once and twice a week of 2 mM Phe and of 4 mM Phe sprayed once a week on the growth of cucumber plants cv. Romi grown in large pots under commercial like conditions and treated once a week for 6 weeks. The evaluated growth parameters were, **a**. plant height (day 43), **b**. nodes number (change between days 5 to 43, **c**. area of leaf 12 (change between days 12 to 19), **d**. number of set fruits (day 50), **e**. number of mature fruits at 56 days, **f**. weight of mature fruits at 56 days, and **g**. weight of single fruits at 56 days. Values in each parameter followed by a different letter are significantly different according to one-way ANOVA with Tukey–Kramer’s HSD test. Default significance levels were set at α = 0.05. Bars = SEs.

Vegetative parameters - Results presented in Table 1

a

Change in plant height (% increase)

|  | change 18 to 39 |
| --- | --- |
| Control | 12.486 |
| 2 mM once/week | 39.2167 |
| 4 mM once/week | 85.7363 |
| 2 mM twice/week | 11.7776 |

b

Change in number of nodes (% increase)

|  | change 18 to 39 |
| --- | --- |
| Control | 23.4343 |
| 2 mM once/week | 39.798 |
| 4 mM once/week | 58.2197 |
| 2 mM twice/week | 41.9066 |

c

Change in leaf length (% increase)

|  | Change 10 vs 3 d |
| --- | --- |
| Control | 119.431 |
| 2 mM once/week | 134.547 |
| 4 mM once/week | 202.902 |
| 2 mM twice/week | 112.544 |

d

Change in leaf length (% increase)

|  | Change 39 vs 27 |
| --- | --- |
| Control | 63.5093 |
| 2 mM once/week | 101.727 |
| 4 mM once/week | 128.01 |
| 2 mM twice/week | 98.1323 |

e

Change in leaflet area (% increase)

|  | Change 10 vs 3 |
| --- | --- |
| Control | 88.3042 |
| 2 mM once/week | 122.253 |
| 4 mM once/week | 281.008 |
| 2 mM twice/week | 125.57 |

Change in leaflet area (% increase)

|  | Change 39 vs 27 |
| --- | --- |
| Control | 98.8876 |
| 2 mM once/week | 99.9759 |
| 4 mM once/week | 148.498 |
| 2 mM twice/week | 117.691 |

g

Plant fresh weight (g)

|  | whole canopy wet |
| --- | --- |
| Control | 372.5 |
| 2 mM once/week | 487.5 |
| 4 mM once/week | 651.25 |
| 2 mM twice/week | 506.25 |

h

Plant dry weight (g)

|  | Dry weight |
| --- | --- |
| Control | 47.505 |
| 2 mM once/week | 64.475 |
| 4 mM once/week | 74.5575 |
| 2 mM twice/week | 66.645 |

Figure S4

Effect of spray once and twice a week of 2 mM Phe and spray of 4 mM Phe once a week on growth parameters of tomato plants cv. Ikram maintained in large pots under commercial like conditions. Phe treatments were carried out until day 73 after the initiation of the applications.

The evaluated vegetative parameters were: **a**. change in plant height from day 18 to day 39; **b**. change in the number of nodes from day 18 to day 39; **c**. change in leaf 5 length from day 3 to day 10; **d**. change in leaf 11 length from day 27 to day 39; **e**. change in terminal leaflet area (at leaf 5) from day 3 to day 10; **f**. change in terminal leaflet area (at leaf 11) from day 27 to day 39; **g**. whole plant fresh weight at day 73; **h**. whole plant dry weight. Values in each parameter followed by a different letter are significantly different according to one-way ANOVA with Tukey–Kramer’s HSD test. Default significance levels were set at α = 0.05. Bars = SEs.

Reproductive parameters - Results presented in Table 1

i

Infloresences (No./plant)

|  | 27 d |
| --- | --- |
| Control | 2.75 |
| 2 mM once/week | 3 |
| 4 mM once/week | 6.5 |
| 2 mM twice/week | 3 |

j

Set fruits (no./plant)

|  | 39 |
| --- | --- |
| Control | 5.25 |
| 2 mM once/week | 6 |
| 4 mM once/week | 8.5 |
| 2 mM twice/week | 4 |

k

Fruits weight (g/plant)

|  | weight |
| --- | --- |
| Control | 679.65 |
| 2 mM once/week | 1034.43 |
| 4 mM once/week | 1512.12 |
| 2 mM twice/week | 678.007 |

l

Fruits (No./plant)

|  | number fruits |
| --- | --- |
| Control | 21.5 |
| 2 mM once/week | 31 |
| 4 mM once/week | 38.25 |
| 2 mM twice/week | 24.75 |

m

Single fruit (g/fruit)

|  | single fruit |
| --- | --- |
| Control | 31.0788 |
| 2 mM once/week | 33.4101 |
| 4 mM once/week | 39.5205 |
| 2 mM twice/week | 27.3956 |

Figure S5

Effect of spray once and twice a week of 2 mM Phe and spray of 4 mM Phe once a week on growth parameters of tomato plants cv. Ikram maintained in large pots under commercial like conditions. Phe treatments were carried out until day 73 after the initiation of the applications.

The evaluated reproductive parameters were **i**. total number of inflorescences in day 27; **j**. total number of set fruits in day 39; **k**. total weight of fruits at day 73; **l**. total number of fruits at day 73; **m**. average weight of a single fruit at day 73. Values in each parameter followed by a different letter are significantly different according to one-way ANOVA with Tukey–Kramer’s HSD test. Default significance levels were set at α = 0.05. Bars = SEs.
